# Expectation and attention increase the integration of top-down and bottom-up signals in perception through different pathways

**DOI:** 10.1101/446948

**Authors:** Noam Gordon, Naotsugu Tsuchiya, Roger Koenig-Robert, Jakob Hohwy

## Abstract

Perception results from the integration of incoming sensory information with pre-existing information available in the brain. In this EEG (electroencephalography) study we utilised the Hierarchical Frequency Tagging method to examine how such integration is modulated by expectation and attention. Using intermodulation (IM) components as a measure of non-linear signal integration, we show in three different experiments that both expectation and attention enhance integration between top-down and bottom-up signals. Based on multispectral phase coherence, we present two direct physiological measures to demonstrate the distinct yet related mechanisms of expectation and attention. Specifically, our results link expectation to the modulation of prediction signals and the integration of top-down and bottom-up information at lower levels of the visual hierarchy. Meanwhile, they link attention to the propagation of ascending signals and the integration of information at higher levels of the visual hierarchy. These results are consistent with the predictive coding account of perception.

## Introduction

Perception is not a simple ‘bottom-up’ mechanism of progressive processing of the sensory input. Instead, perception is made possible by processing sensory information in the context of existing information in the brain spanning multiple levels of the cortex. The ability of the visual system, for example, to reach unambiguous representations from highly complex, variable and inherently ambiguous sensory inputs can be understood in terms of Bayesian inference and probabilistic integration of prior knowledge (top-down) with stimulus features (bottom-up). Crucially, within these kinds of framework, the integration of top-down and bottom-up signals is dynamically modulated by cognitive and potentially interacting cognitive factors such as expectation and attention (1–5).

While expectation and attention are much studied, attempts to dissociate the two and study their unique yet interrelated underlying mechanisms are relatively recent and far from complete (6–10). Great caution in experimental design is required in order to obtain empirical data that permit a genuine distinction between the neural processes underlying expectation and attention. A fundamental challenge is to keep expectation and attention sufficiently separate; studies tend to either rely on explicit probability cues, which introduces task demands, or to vary stimulus properties across conditions, which confounds claims to have revealed high-level effects. The paradigms we introduce in this study were designed specifically to avoid such pitfalls.

To consider the potentially distinct roles of expectation and attention we here appeal to the predictive coding theory of perception. In predictive coding, the brain’s ability to infer the causes of its sensations is attributed to its ability to embody the statistical structure in the environment within a generative model describing the hierarchical and dynamic statistics of the external world (2, 11). Perception is understood to reflect the process of inferring the causes and states in the external world that cause the sensory input. Expectations, under this framework, allow predictive signals to descend from higher to lower levels in the cortical hierarchy, where they are tested against sensory-driven information. The gap between the two - the prediction error - gives rise to the ascending signals which, in turn, allow for the higher-level expectations and subsequent predictions to be optimised in an iterative fashion. The outcome of this hierarchical prediction error minimization (resolution) process is, according to predictive coding, perception.

Prediction errors may result from two related sources: inaccurate top-down predictions (that do not match the actual state of the external environment), or imprecise or noisy bottom-up sensory information (such as vision in a foggy day or hearing through a brick wall). An efficient system should therefore incorporate an estimation of the precision (i.e., inverse of variance) of sensory signals. The more precise a prediction error is estimated to be, the more ‘reliable’ it is considered to be, leading to more revision of the generative model. Attention has been suggested to reflect such a process of optimizing precision estimates of prediction errors, which, in more mechanistic terms, may allow the ‘prioritisation’ of signals expected to be more precise by means of increasing the synaptic gain of neuronal units encoding precision estimates (Figure 1A) (3). Expectation and attention therefore relate, under the predictive coding framework, to descending and ascending signals, respectively.

**Figure 1.**
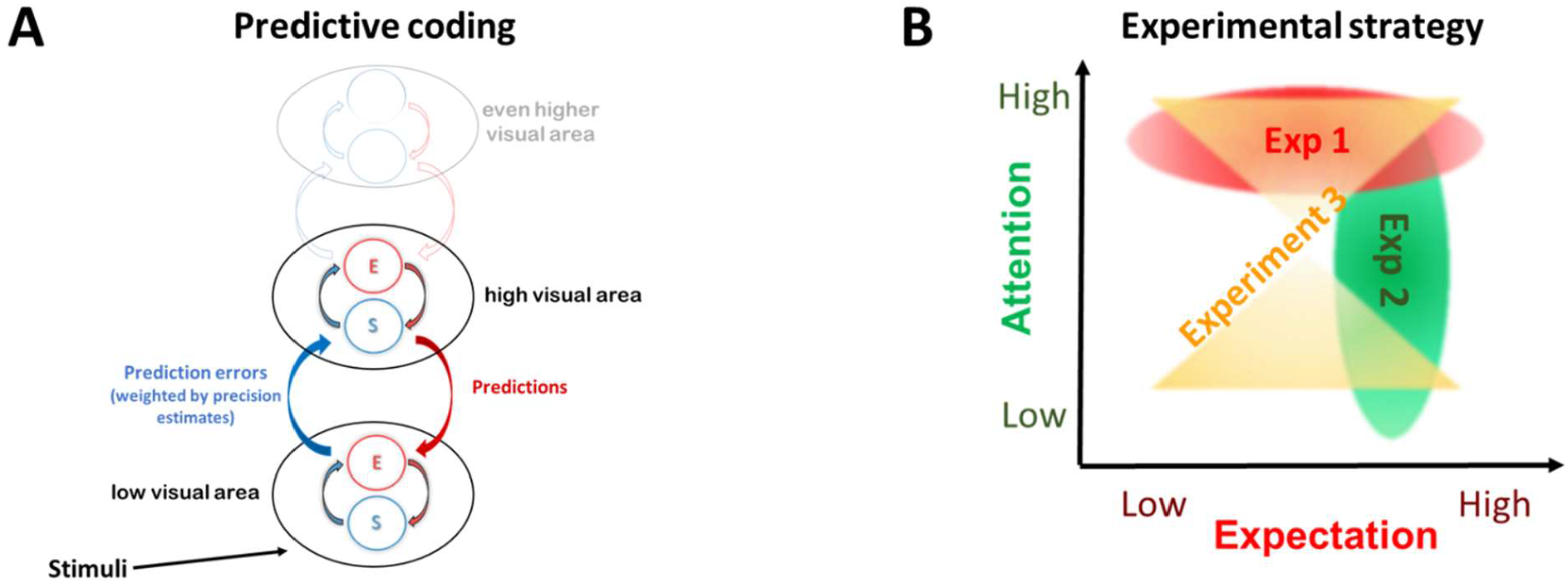
Theoretical background and experimental strategy. (A) The predictive coding theory of perception, which describes perception as the inference made about the state of the external world and the causes of the sensory input. Under this framework, expectations allow predictive signals to descend from higher to lower levels in the cortical hierarchy, where they are tested against sensory-driven information. The discrepancy between the two - the prediction error - propagates up the hierarchy, allowing for the higher-level expectations and subsequent predictions to be optimised. Expectations and prediction errors are suggested to be coded within each hierarchical level by distinct neural populations referred to as state (S) and error (E) units, respectively (14, 15). Prediction errors are suggested to be weighted by their estimated (and expected) precision such that high precision estimates lead to enhancement of prediction error signals via synaptic gain mechanisms. (B) Conceptual figure of the 3 experiments used to comprehensively examine the role of expectation and attention in the integration of top-down and bottom-up signals. In experiment 1 (red oval) expectation was manipulated while endogenous attention and the visual stimuli were kept constant across conditions. In Experiment 2, attention was manipulated while expectation and the visual stimuli were held constant across conditions (green oval). In experiment 3, a novel analysis method of data obtained in a previous study is used to simultaneously examine main effects of expectation and attention, as well as the interaction between the two (yellow hourglass shape). For consistency, all figures in Results use shades of red for expectation, and shades of green for attention.

Several implications follow from this account of expectations, predictions, prediction error minimization and attention. On the one hand, highly predictable stimuli are expected to yield smaller prediction errors and thus attenuated prediction error-related neural activity. On the other hand, if attention estimates high precision of the signal (i.e., greater prediction error impact), then even expected stimuli will yield greater prediction error-related neural activity. Indeed, various studies suggest an interaction between attention and expectation, and have demonstrated that when stimuli are unattended (e.g., they are task irrelevant) high levels of expectation can result in reduced sensory signals; however, when stimuli are attended (e.g., they are task-relevant) expected stimuli can, in fact, result in greater neural activity ((12) but also see Garrido *et al.* (13)).

The goal of the present study is to elucidate the mechanisms underlying attention and expectation and to better understand the relationship between these factors in perception. To do so, we performed three experiments and analyses aimed at comprehensively studying these factors while avoiding the potential pitfalls described above (Figure 1B).

All experiments utilized the Hierarchical Frequency Tagging (HFT) method in EEG ((16); Figure 2A and Videos 1 and 2). The HFT method was designed to investigate hierarchical visual processing. Its strength lies in its ability to distinguish between neural signals derived from different cortical levels, while providing a measure for the integration of these signals. In brief, two frequency-tagging methods are combined – the Steady-State Visual Evoked Potentials (SSVEP; (17, 18) and the Semantic Wavelet-Induced Frequency-Tagging (SWIFT; (19). While SSVEP originates primarily in lower visual areas in the occipital cortex (20), SWIFT has been shown to selectively tag high-level object representation areas but not early visual areas in both EEG (19) and fMRI (21). Spectral power of the recorded signal at the tagged frequencies for the SSVEP and SWIFT are understood to reflect lower- and higher-level cortical activity, respectively.

**Figure 2.**
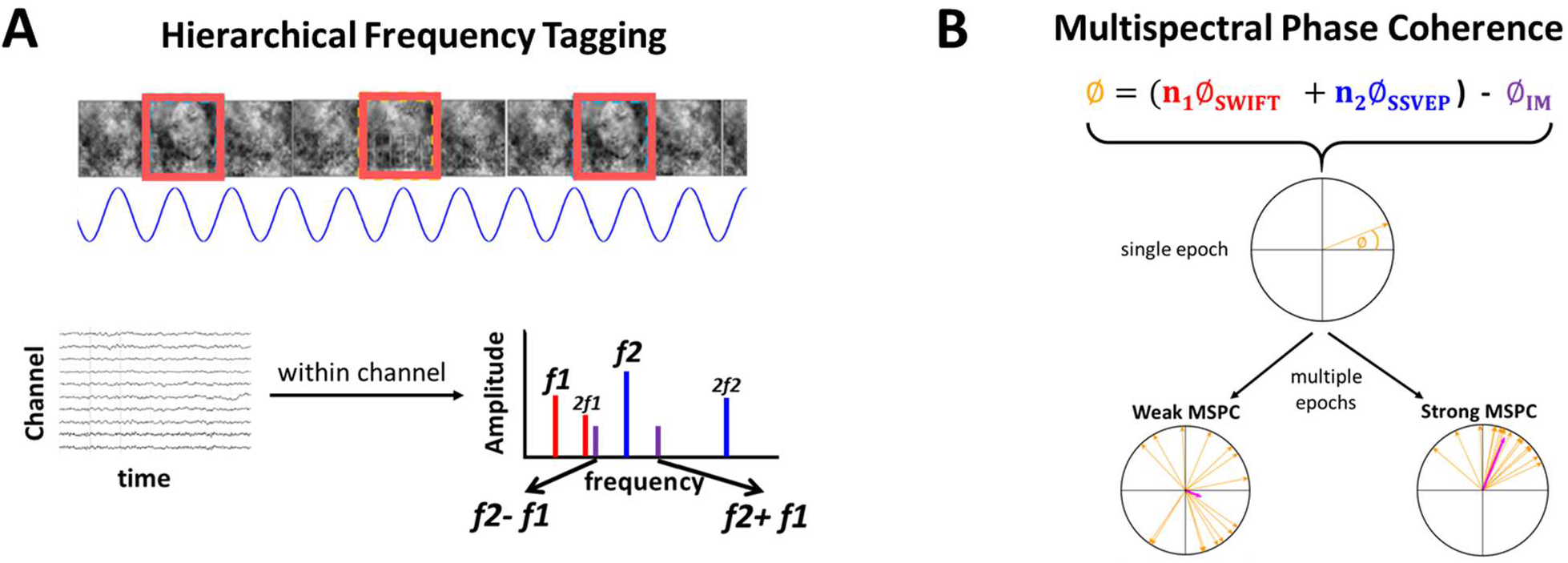
Stimulus construction and analysis methodologies. (A) All experiments implemented the Hierarchical Frequency Tagging (HFT) method using face and house images. Semantic Wavelet-Induced Frequency-Tagging (SWIFT) sequences are presented at given frequencies, allowing tagging of image-recognition activity (red rectangles). Contrast-modulation is applied at a higher frequency inducing SSVEP (blue sinusoid). When analysing the EEG data in the frequency domain (bottom graph with multiple peaks), peaks in the power spectrum can be seen at the fundamental frequencies and their harmonics (red bars for SWIFT f1 and blue bars for SSVEP f2). Additional peaks at IM components (e.g. purple bars for f2+f1 and f2-f1) are suggested to indicate integration of bottom-up SSVEP-driven signals with top-down SWIFT-driven signals. (B) The Multi Spectral Phase Coherency (MSPC) (28) quantifies the degree to which an IM frequency component is driven by the phases of the fundamental input frequencies. In other words, the degree to which the IM component reflects an *interaction* between those input frequencies. Within each epoch, we first calculate the difference between the sum of the (weighted) phases of the fundamental input frequencies, and the phase of the IM component. Then, we compute the coherence of this value across multiple epochs applying the same method as in the well-known phase-locking value (see Methods for a detailed description.) Here, we introduced a novel distinction between two measures – MSPCstim and MSPCres - which differ in what they consider the ‘input’ signals to be. Specifically, the MSPCstim ties the IM phase to the phases of the stimulus itself (i.e. the images presented on the screen), while MSPCres ties it to the phases of the tagged neural responses (Figure 7). We suggest that these measures distinguish between neural interactions occurring at lower and higher cortical levels, respectively.

Crucially, intermodulation (IM) components, that is, linear combinations of the fundamental input frequencies, serve as a measure of non-linear interaction. In the brain, non-linear neural interactions enable rich, context-dependent information processing and play a key role in perception (22–24). Indeed, several EEG studies have utilized IMs to reveal the mechanisms of visual object-recognition (25–27). In the case of HFT, intermodulation of the SWIFT and SSVEP frequencies are understood to reflect integration of top-down SWIFT-driven signals with bottom-up SSVEP-driven signals.

IMs are typically quantified using amplitude-based measures. Importantly, we introduce a novel distinction between two phase measures of the IM signal, based on the Multi Spectral Phase Coherence (MSPC; (28)). The first measure, *MSPCstim*, ties the IMs to the phases of the SWIFT and SSVEP *stimulus* modulation, and the second one, *MSPCres*, to the tagged SWIFT and SSVEP neural *response*. We argue these two measures distinguish neural signal integration occurring at different hierarchical levels (Figure 2B, and described in greater detail in Discussion and Methods). We hypothesised that if expectation and attention indeed relate to descending and ascending signals (respectively), their influence should manifest differently in these two measures.

Two new paradigms were designed to ensure two critical experimental aspects: that expectation and attention are manipulated individually without confounding each other, and that their modulatory effects are determined cognitively without introducing demand characteristics and without coinciding with any differences in the visual stimuli. In addition to these, new analyses performed on previously published data (Gordon *et al.*, 2017) allowed a direct examination not only of expectation and attention main effects but of the interaction between the two. Indeed, the two phase measures we introduce here were modulated differently by expectation and attention, providing direct physiological evidence for their distinct hierarchical modulation of perceptual processing.

## Results

### Expectation-Experiment 1

#### Behavioral task

Experiment 1 manipulated expectation by using two tasks, specifically designed to avoid explicit indication of expectation levels in the instructions (thus minimizing demand characteristic), while holding attention and the visual stimuli constant across conditions. In each HFT trial a series of consecutive SWIFT-scrambled sequences (of one house and one face image) were presented to participants (Video 3) who were required to perform one of two tasks. In the image-repetition (IR) task, participants were requested to press the space-bar when either image repeated itself either 3 or 4 times (as instructed before each trial). Stimuli in these trials were considered ‘unexpected’ as participants could not predict the upcoming image. In the pattern-violation (PV) tasks, participants were required to memorise a pattern of 5-6 images presented to them as text (‘Face, House,…’ etc.) before the trial. When the trial began, the pattern repeated itself over and over and participants were instructed to press the space-bar when the pattern was violated. Thus, in PV trials, almost all upcoming images were highly predictable thus expectations were much more reliable than in the IR trials. Each series of face and house images (e.g. FHHFHFHHFH…) appeared in one PV and in one IR trial (using different images), such that the series used for both task were, in fact, identical.

Participants were tested for 4 blocks, in the order of PV-, IR-, PV- and IR- block. After completing all 4 blocks, participants were asked to compare between the PV and the IR tasks and to report whether they noticed the underlying patterns in the IR tasks. Indeed, despite the PV blocks preceding their matching IR blocks (see Methods), only N=3 out of 15 confirmed noticing an underlying pattern in some IR trials. This validates our assumption that the PV and IR tasks manipulated the expectation for upcoming stimuli. Second, N=10 out of 15 participants reported finding it more difficult to recognise the actual images of the IR compared to the PV trials. Several participants reported that the house and the face images tended to perceptually ‘blend’ more with each other in the IR trials (note that some elements of the images, such as edges with strong contrasts, may remain visible to some degree also in the ‘scrambled’ frames, accounting for why images could sometimes be perceived as blended; see Video 3). Given that the same method was used to construct all stimuli in the experiment, the reported differences in perception can be strictly attributed to the task instructions, highlighting the impact of one’s expectation on conscious perception.

### EEG analysis-Expectation modulates MSPCstim but not MSPCres

After applying the fast Fourier Transform (FFT) on the EEG data of each trial, we verified that tagging was obtained for both the SWIFT and the SSVEP frequencies (SWIFT = f1 = 1.2 Hz, and SSVEP = f2 = 15 Hz). Peak amplitude SNRs at both fundamental frequencies and their harmonics were evident in the FFT spectrum averaged across all electrodes, trials, participants (Figure S1A).

Critically for the purpose of this paper, we examined the effect of expectation (predictability) on the IM signals. Based on the Multispectral Phase Coherency (MSPC), we quantified the degree to which the IM phases were driven by the SWIFT and SSVEP phases. As detailed in Methods, we introduce a novel distinction between the MSPCstim measure in which the *stimulus* (image) SWIFT and SSVEP phases are considered the driving inputs, and the MSPCres measure in which the EEG SWIFT and SSVEP neural *response* phases are considered the driving inputs (both MSPCstim and MSPCres were computed individually within each electrode). As detailed in Discussion, we suggest these measures to indicate cortical signal integration occurring at different levels.

MSPC values were calculated individually for each channel (with reference to its own SSVEP and SWIFT phases) within each trial (see Methods). Analyses were then performed on the average of both second-order IM components (f2-f1=13.8Hz and f2+f1=16.2Hz) in a posterior ROI (17 electrodes) including all occipital (Oz, O1 and O2), parieto-occipital (POz, PO3-PO4, and PO7-PO8) and parietal (Pz and P1-P8) electrodes. As shown in Figure 3, MSPCstim was higher for the PV (expected) trials compared to the IR (unexpected) trials (χ2 =22.9, p<0.001), indicating increased neural integration between the SWIFT and SSVEP signals when stimuli are expected. This effect was not evident for the MSPCres measure (χ2 =1.36, p>0.05). The significance of the MSPCstim vs. MSPCres result will be discussed later when comparing the relationship between these measures across all experiments and analyses.

**Figure 3.**
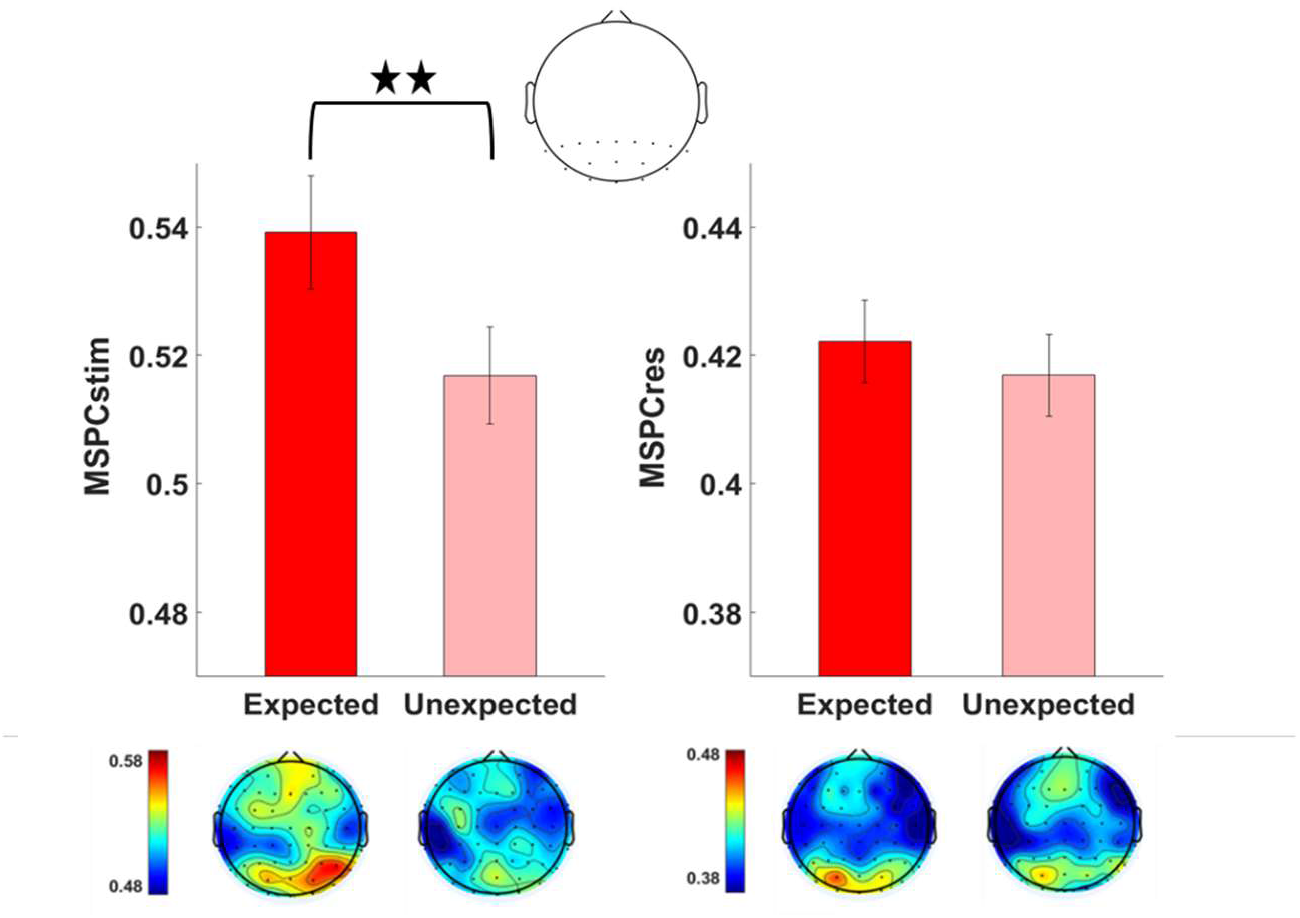
Expectation modulates MSPCstim but not MSPCres. Multi Spectral Phase Coherence (MSPC) averaged across the two second order IM components (f1+f2 and f1-f2) in the expected and the unexpected conditions. Results are shown for a posterior ROI (17 electrodes, top) and the scalp topographies (bottom). Error bars represent standard error across subjects (N=15). The MSPCstim measure (left) quantifies IM responses by examining the degree to which the IM phase is driven by the phases of SWIFT and SSVEP stimulus (image) modulation. Conversely, the MSPCres measure (right) examines the degree to which the IM phase is driven by the tagged SWIFT and SSVEP neural response phases. These measures are therefore suggested to indicate signal integration occurring at earlier and at later stages of cortical processing, respectively. MSPCstim (right) was higher for the PV (expected) trials compared to the IR (unexpected) trials (χ2 =22.9, p<0.001), indicating increased neural integration between the SWIFT and SSVEP signals when stimuli are expected. This effect was not evident for the MSPCres measure (χ2 =1.36, p>0.05).

### Attention-Experiment 2

#### Behavioral task

In experiment 2, HFT trials with house and face images were presented to participants with contrast modulation (SSVEP) at 12Hz. Unlike experiment 1, two different SWIFT frequencies were used for the house and the face images (0.8Hz and 1Hz, counterbalanced across trials). Importantly, images were superimposed via alpha-blending, enabling simultaneous tagging of both frequencies (one for each image type) within each trial. In each trial, participants were requested to count either the faces or the houses, or to perform a demanding central-attention task. These behavioral tasks defined each image as either ‘attended’ or ‘unattended’ within each trial. Note that only 70-85% of the SWIFT cycles contained the face or house images while the rest of the cycles contained their matching ‘noise’ sequences (See Methods). This ensured that the attentional task was sufficiently demanding (Video 4).

#### EEG analysis-Attention modulates MSPCres but not MSPCstim

First, we verified that we were able to obtain separate tagging for the relevant frequencies (two SWIFT: 0.8Hz and 1Hz, and SSVEP: 12 Hz) (Figure S1B). Indeed, peak amplitude SNRs at all three fundamental frequencies were evident in the FFT spectrum averaged across all electrodes, trials and participants (Figure S1B).

As in experiment 1, MSPC values were calculated individually for each channel within each trial (see Methods) and statistical analyses were performed on the average of both second-order IM components in a posterior ROI. Interestingly, as opposed to results from experiment 1, the effect of attention was evident in the MSPCres measure but not MSPCstim (Figure 4). Specifically, MSPCSres was higher for the attended compared to the unattended images (χ2 = 41.4, p≪0.001). This effect was not evident for the MSPCstim measure (χ2 = 1.21, p>0.05).

**Figure 4.**
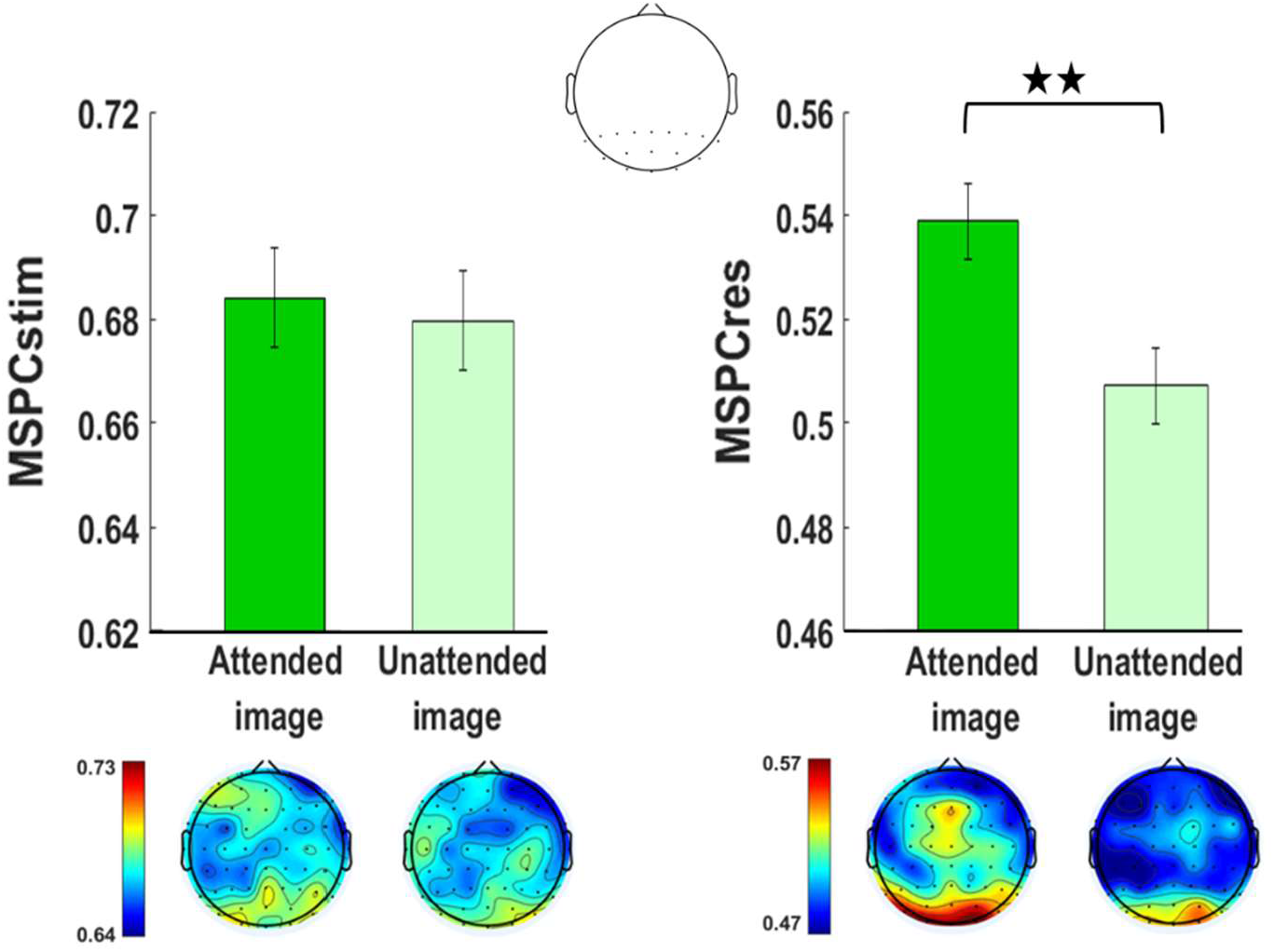
Attention modulates MSPCres but not MSPCstim. Multi Spectral Phase Coherence (MSPC) averaged across the two second order IM components (f1+f2 and f1-f2) in the expected and the unexpected conditions. Results are shown for a posterior ROI (17 electrodes, top) and the scalp topographies (bottom). Error bars represent standard error across subjects (N=11). The MSPCstim measure (left) quantifies IM responses by examining the degree to which the IM phase is driven by the phases of SWIFT and SSVEP stimulus (image) modulation. Conversely, the MSPCres measure (right) examines the degree to which the IM phase is driven by the tagged SWIFT and SSVEP neural response phases. These measures are therefore suggested to indicate signal integration occurring at earlier and at later stages of cortical processing, respectively. MSPCres (left) was higher for the attended compared to the unattended images (χ2 = 41.4, p≪0.001), indicating increased neural integration between the SWIFT and SSVEP signals when stimuli are attended. This effect was not evident for the MSPCstim measure (χ2 = 1.21, p>0.05).

Further analysis of the data from experiment 1 and 2 show comparable results for various higher order IMs, and is consistent with the suggestion that the attentional modulation manifests at higher cortical levels (Supplemental Information). In brief, the effect of attention was most notable for the 4^th^ order IMs (2f2±2f1) and additional analysis suggests this effect to reflect interactions occurring at later rather than earlier processing stages.

### Interaction of attention and expectation-reanalysis of previous study

After differentiating expectation and attention in Experiments 1 and 2, and establishing that both expectation and attention are associated with enhanced IMs, we returned to our previously published data (16) to examine the interaction between these factors, and to evaluate the consistency of the results from that study with those of experiment 1 and experiment 2 here (Supplemental Information). The interaction between expectation and attention was not significant for MSPCstim (χ2 = 3.47, p>0.05) but it was indeed highly significant for MSPCres (χ2 = 19.56, p<0.001). In fact, the slope of MSPCres against expectation was negative for unattended images (χ2 = 5.05, p<0.05). Additional post-hoc analyses performed individually for expectation and attention were consistent with the results from experiment 1 and Experiment 2, where MSPCstim showed greater enhancement with expectation and MSPCres showed greater enhancement with attention (Supplemental Information). These results are interpreted further in Discussion.

Finally, we performed several analyses to examine the relation between the MSPC and amplitude measures. Specifically, we examined whether reduced SWIFT, SSVEP and/or IM response amplitudes can account for the reduction of MSPCres in the unattended compared to the attended conditions. While we expect the MSPC and amplitude measures to correlate to some degree, we confirmed that the amplitude measures accounted for no more than 20% of the MSPC variance. Further details of the analyses are provided in Supplemental Information.

## 4. Discussion

The goal of this study was to examine the mechanisms underlying attention and expectation in perception, focusing on their modulation of top-down and bottom-up signal integration. All experiments utilised the Hierarchical Frequency Tagging (HFT) method (16) in which SSVEP- and SWIFT- driven signals reflect activity in lower (V1/V2) and higher levels in the visual hierarchy, respectively (19, 21). We reasoned, within the predictive coding framework, that the SWIFT-tagged signals reflect top-down, semantically rich predictions while SSVEP-tagged signals reflect bottom-up sensory input (or prediction errors). Importantly, intermodulation (IM) frequency components are a distinct and objective indicator for non-linear integration of multiple input frequencies. Accordingly, IMs are hypothesised here to be influenced by, and provide an indication of, the level of integration between top-down predictions and bottom-up prediction-error signals.

Two new experiments and a reanalysis of an existing data set were examined in this study, covering a range of experimental modulations of expectation and attention. The first experiment modulated expectation while holding attention constant, and the second experiment modulated attention while holding expectation levels constant. Critically, both experiments used the same sensory input across conditions, while cognitive modulations were achieved only by means of the behavioral tasks at hand. Our results provide direct neural evidence for the increased integration of bottom-up and top-down signals through modulation of expectation and attention.

Importantly, we argue that the dissociation between the MSPCstim and MSPCres measures found here relates expectation and attention to distinct mechanisms impacting the integration of descending and ascending signals at lower and at higher levels of the visual hierarchy, respectively.

We believe this is the first direct demonstration of the different neurophysiological manifestations of the mechanisms by which expectation and attention change the very integration of top-down signals with bottom-up signals in perception.

In the following, we show how our results can be both explained by, and provide support to the predictive coding model of perception (Figure 1A).

### 4.1 Terminology and theoretical background

The terms expectation and attention have been confounded in various cognitive studies (for discussion, see (8)). Furthermore, both attention and expectation are composed of many different aspects (e.g. spatial vs. feature-based attention, selective vs. sustained attention, etc.; expectation can result from implicit learning of statistical contingencies or through explicit instruction, etc.), which can have distinct neural origins and underlying mechanisms. Like Summerfield et al. (8), we use the term attention to denote the motivational relevance of an event and the term expectation to denote its conditional probability.

The terms ‘predictions’ and ‘expectations’ can be used in the more technical predictive coding literature to denote somewhat different aspects of the perceptual circuitry. Predictions are the descending signals that are used to explain away sensory or neural input in lower areas. Expectations on the other hand are the inferences made about the state of the external world (or more technically speaking, the inferred values assigned by the generative model to the hidden causes and states in the external world). In that sense, predictions are compared against sampled sensory inputs in the sensory cortex or against expectations at intermediate hierarchical levels (3, 14). We use ‘expected’ and ‘predictable’ interchangeably here.

Attention, in cognitive neuroscience literature, is commonly viewed as mechanisms which: 1) increase baseline neural activity, sometimes referred to arousal or alertness, 2) selectively enhance relevant neural responses, and 3) selectively inhibit irrelevant responses (29). In predictive coding, such signal modulation is suggested to be achieved by means of internal estimates (and expectations) of precision (the inverse of signal variability). Greater precision estimates lead to greater weighting of the prediction error, effectively serving as a gain control for the bottom-up signals and allowing greater modification of higher-level expectations and predictions (30, 31). Attention, under this framework, enables signal enhancement by modulating the gain on the prediction errors obtained through varied estimations of prediction-error precision estimations (3).

EEG signals are thought to be generated by the spread of postsynaptic potentials along the apical dendrites of pyramidal cells (32). In other words, they are generated by activity at afferent rather than efferent pathways. With this in mind, the modulatory effects of expectation and attention, as described above, can be expected to manifest differently at lower and higher visual areas, respectively.

### 4.2 What can be learnt from the intermodulation components

As the IM components are key to our analysis, it is important to clarify several issues. First, the IM signal depends on two main factors: the amplitude and phase of the input signals, and the specific mechanism of signal integration at hand. The former factor is straightforward as changing the power or shifting the phase of the input can lead to a shift in output power or phase, respectively. Using IMs to infer something about the latter factor is, however, less trivial since non-linear neuronal dynamics may be consistent with various models of neural processing, ranging from cascades of non-linear forward filters (e.g., convolution networks used in deep learning) through to the recurrent architectures implied by predictive coding. It is therefore not easy to link specific computational or neuronal processes to the IM responses and IMs can only provide indirect evidence for predictive coding as well as other theories of perception. However, as we will argue, various arguments indeed point to the recurrent and top-down mediation of the IM responses in our data.

The second issue to consider is the interpretation of the MSPC measure in the context of our novel distinction between MSPCstim and MSPCres. As described in Methods, the MSPC measure aims to quantify the degree to which the phase of an IM component is driven by the phases of the input frequencies. The question then is what we consider those driving inputs to be? SSVEP and SWIFT phases can be quantified either by the stimulus itself or by the EEG response signal (as tagged and obtained by the FFT). We term these options MSPCstim and MSPCres, respectively. When considering these two measures, one may *a priori* expect them to behave quite similarly. After all, the stimulus (image) and the response (EEG) phases can be expected to be highly correlated with each other, leading to highly correlated MSPCstim and MSPCres measures. Threfere, our finding here is highly non-trivial; the expectation and attention manipulations influenced these measures in a highly different manner (Figures 3-5), directly indicating distinct neural processes for expectation and attention. As we describe in greater detail in Methods, we reason that MSPCstim and MSPCres may allow a distinction between interactions occurring at lower and higher levels of hierarchical processing, respectively. Hence, while MSPCstim and MSPCres may be correlated, distinct underlying correlates should account for the differences in results obtained across the three experiments.

**Figure 5.**
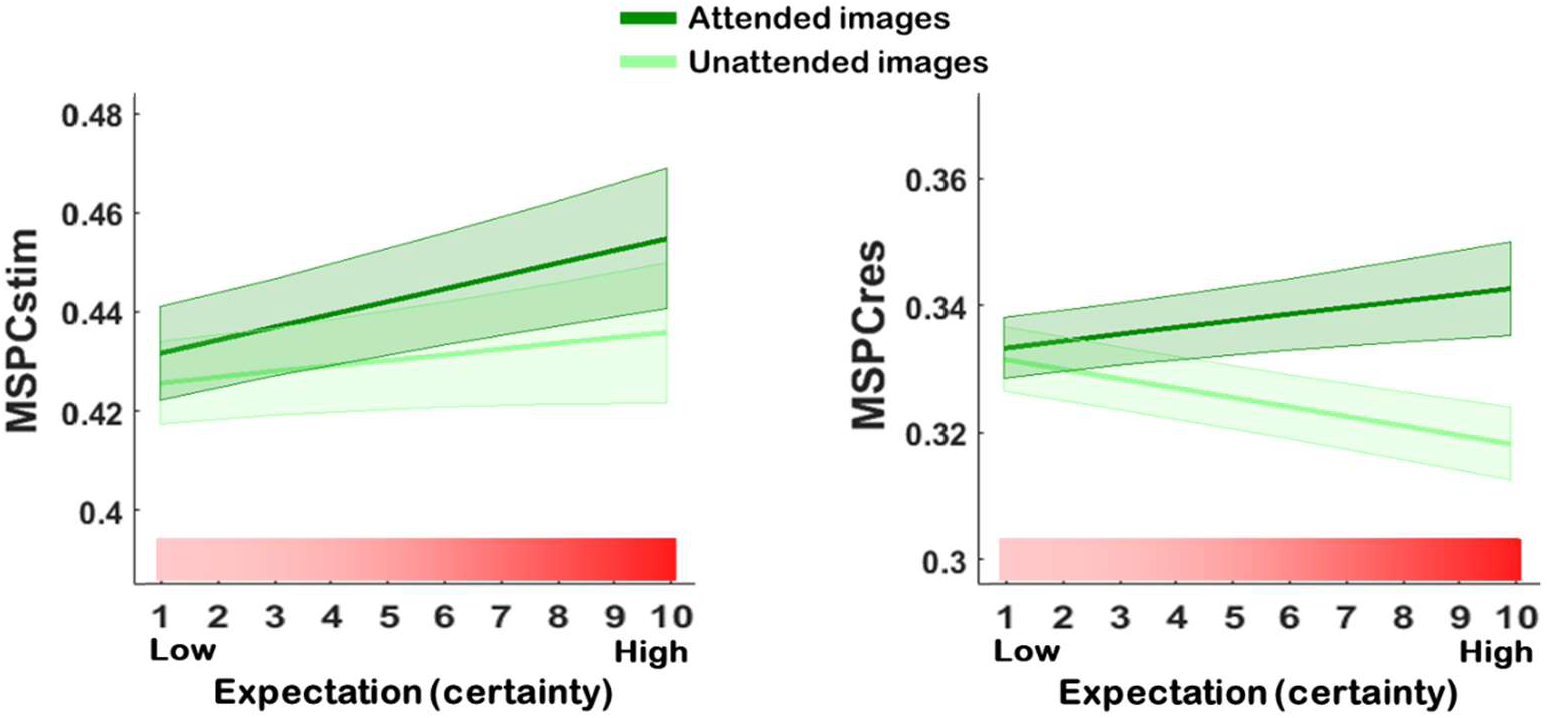
The expectation-attention interaction is significant for MSPCres but not MSPCstim. Predicted MSPCstim (left) and MSPCres (right) values obtained from a full Linear Mixed Effects interaction model, with their standard error indicated by the shaded area. The model included expectation, attention and an expectation-attention interaction term as the fixed effects, while the random effects included a random intercept for frequency nested within channels nested within participants, and random expectation and attention slopes for each participant. Consistent with the colors used in the previous figures, attended images are represented by the dark green lines and unattended images by the light green lines, while the pink-red gradient indicates increasing expectation. The significance of the interaction term was tested using the likelihood ratio test between the full model and the reduced model which excluded the interaction fixed effect. The expectation-attention interaction was not significant for MSPCstim (χ2 = 3.47, p<0.05) but was highly significant for MSPCres (χ2 = 19.56, p<0.001).

### 4.3 Expectation and attention: main effects and their interaction

Combining results from all experiments, we suggest MSPCstim and MSPCres to be more strongly related to the expectation, and to the attention modulation, respectively.

The MSPCstim measure demonstrated a consistent increase with expectation, as evident in the data obtained from experiment 1 and from Gordon *et al*. (16). Describing a stimulus as being predictable implies that the prediction signal precedes the onset of the stimulus itself. Hence, when the sensory input arrives, the prediction can be tested against (interact with) the incoming sensory-driven information in a highly ‘online’ manner. In such conditions, as in the PV trials in experiment 1 and the ‘high-certainty’ trials in Gordon *et al.*, top-down predictions and bottom up sensory evidence can interact quickly at early visual areas, and the resulting IM phase can be expected to be strongly related to the stimulus phase (see suggested primary source of MSPCstim in Figure 6). This observation is consistent with various studies demonstrating effects of expectation at early visual areas (33–35).

**Figure 6.**
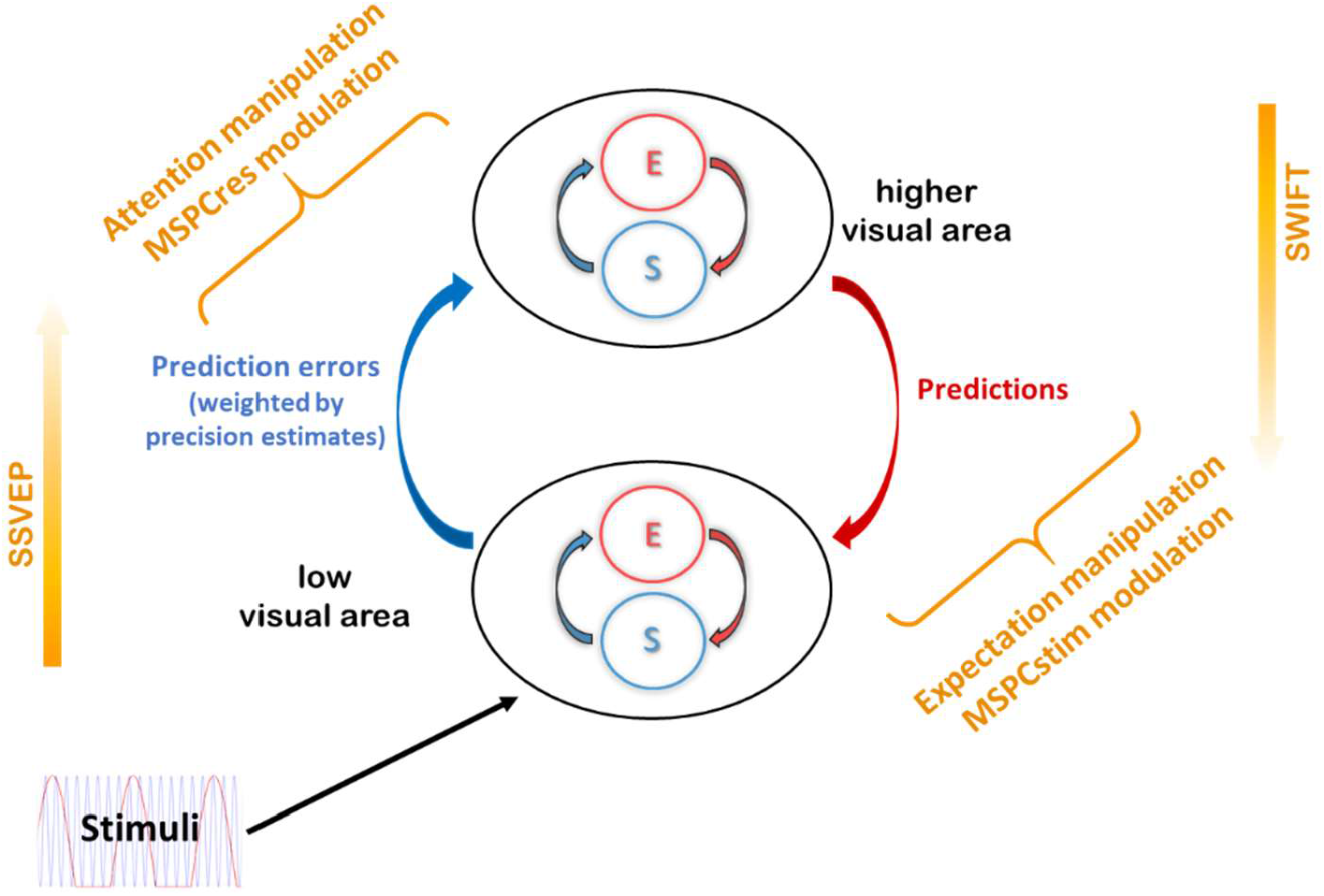
Expectation, attention, MSPC and predictive coding. Results presented in this study can be accounted for by the predictive coding framework of perception as follows: 1) Expectation (the probability for the appearance of specific stimuli) relates to descending prediction signals. 2) Better predictions (as afforded by the PV trials in experiment 1) increase the efficiency of top-down and bottom-up signal integration at low-level visual areas, as reflected by the increased MSPCstim with expectation (Figures 2 and 4). 3) Attention reflects a (precision-weighted) control mechanism for the propagation of prediction error-signals. 4) Attention effectively increases the influence of prediction error signals on expectations at higher hierarchical levels, as reflected by the increased MSPCres with attention (Figures 3 and 4). 5) The effect of expectation on the integration of top-down and bottom-up information at lower visual areas is less dependent on attention than the integration at higher levels. Hence, while MSPCstim increased with expectation for both attended and unattended stimuli (Figure 5), the influence of expectation on MSPCres was attention-dependant (Figure 5).

**Figure 7.**
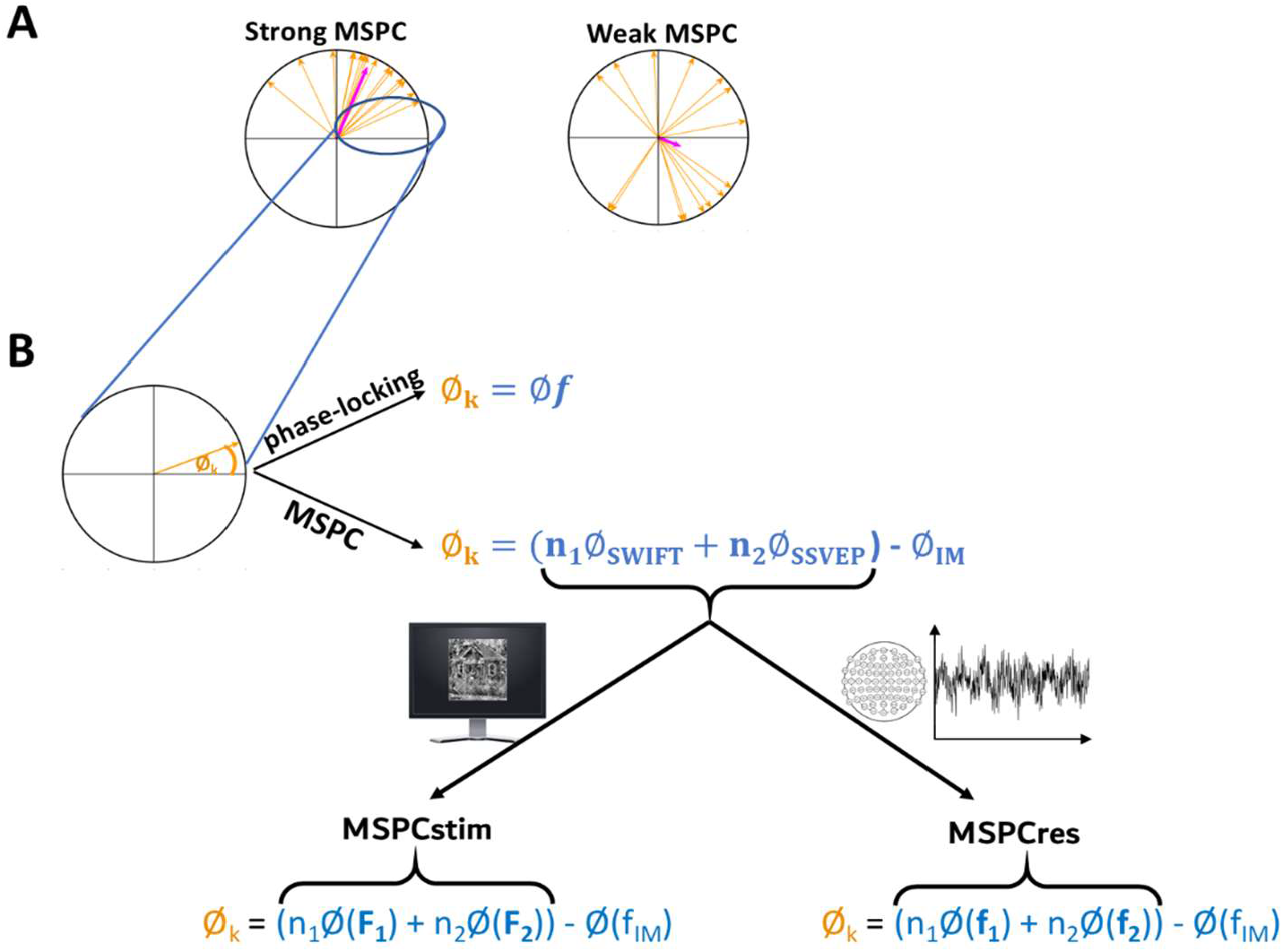
Multi Spectral Phase Coherence. The method for calculating MSPC. (A) Schematic example of weaker (left) and stronger (right) phase coherence. Both the phase-locking value (PLV) and the MSPC measures examine the consistency of a given phase term across multiple epochs. This can be visualised by first converting the phase term from each epoch into a unit (length=1) vector pointing at its phase angle. Phase coherence is then obtained by computing the average vector (the sum of all vectors divided by the number of epochs). The result is a vector whose length can vary from 0 (each vector pointing at random directions, no phase coherence across epochs) to 1 (all vectors pointing at the same direction, perfect coherence across epochs). (B) The primary difference between PLV and MSPC measures is the phase-term used for each epoch. For the PLV, only the phase of one specific frequency (or frequency band) is extracted for each channel/epoch and is used as the phase-term for computing phase-locking. When examining coherence between distant channels, the phase-term used for the PLV would be the difference (Δϕ) between the phases extracted from the different channels. In contrast, the phase-term used for the MSPC calculations here is based on the difference (Δϕ) between the phase of a specific IM component and the (weighted) phases of the fundamental frequencies *within each channel/epoch*. In other words, instead of examining the extent to which the phase is driven by an event in time, the MSPC examines the extent to which the IM phases are driven by the phases of the fundamental frequencies. Our distinction between MSPCstim and MSPCres is reflected by the two MSPC formulas shown at the bottom of the figure. Specifically, MSPCstim defines the phases of the *stimuli* (the on-screen image) as the driving fundamental frequencies (left formula, upper case F1 and F2), and MSPCres defines the *EEG response* phases as the driving fundamental frequencies (right, lower case f1 and f2). Note, that the weights of the fundamental frequencies in those formulas (n_1_ and n_2_) are the coefficients that define the IM frequency (e.g., given F1= 1.2 Hz and F2= 15 Hz, the weights for the 3^rd^ order IM component 2*F1 + F2= 17.4 Hz would be n_1_=2 and n_2_=1).

The MSPCres measure, on the other hand, showed a greater increase with attention in experiment 2 and the Gordon *et al.* data compared to the MSPCstim, with no consistent modulation by expectation. Here, we have suggested MSPCres to be more strongly related to signal integration occurring at higher levels of the visual hierarchy. Attention, in predictive coding, is suggested to optimise perception by allowing the ascending error signals to exert a greater influence on the expectations at the higher cortical level (31). Therefore, attended (compared to unattended) images should allow greater integration of bottom-up error signals with higher level expectations, as reflected by the MSPCres (see suggested primary source of MSPCres in Figure 6). Additional support for this claim is provided by the MSPCres analysis of the 4th-order IMs, which suggests the modulatory effect of attention to involve interactions occurring a later stage than initial sensory processing (see SI).

In addition, an interesting relationship between expectation and attention follows the account of precision-weighted prediction errors (Figure 1A). On the one hand, highly predictable stimuli will yield small prediction errors. On the other hand, if a stimulus is highly relevant (attended), the influence of the prediction-errors on the higher-level expectations will be enhanced (31). Indeed, the significant interaction found for the MSPCres measure (Figure 5, right panel) supports such a relationship between expectation and attention. When stimuli were unattended, MSPCres decreased with higher expectation, in line with reduced prediction error signals. However, when stimuli were attended, not only did MSPCres not decline with expectation, it in fact trended upward.

Given that EEG signals are understood to be generated primarily by activity at afferent synapses, and in accordance with current predictive coding models, we argue that all our results can be accounted for by the association of predictions with descending signals and attention with ascending, precision-weighted error signals (as illustrated in Figure 6). When stimuli are expected, predictions become more accurate, and their integration with sensory evidence at low visual areas improve. Hence, the MSPCstim response, which we suggest reflects neural integration at early visual areas, is enhanced with expectation, regardless of attention. In turn, when stimuli are attended, the propagation (or influence) of ascending error signals to (or on) the higher hierarchical level is enhanced. Hence, the MSPCres response, which we suggest reflects neural integration at later visual areas, is enhanced with attention. In contrast, when the expected stimuli are unattended (not task-relevant), not only are the prediction-errors gated out by (un)attention, there are also smaller to begin with, hence the reduced MSPCres.

While we choose to interpret our results under the predictive coding framework of perception, alternative explanations should be considered as well. One possibility is that some of the differences between the MSPCstim and MSPCres responses may be attributed to the locations of the neural generator sources mediating our expectation and attention modulations. For example, the posterior ROI used in our analyses may have captured attention-related parietal activity more so than expectation-related frontal activity.

Whether and how expectation and attention interact in perception is an ongoing debate in the scientific literature. Several studies have demonstrated a relationship between expectation and attention consistent with the interaction reported here. For example, using multivariate pattern analysis in fMRI, Jiang *et al.* demonstrated that the ability to discriminate expected and unexpected stimuli was notably enhanced with attention (36). In a different fMRI study, Kok *et al.* showed that a significantly reduced neural response to predicted stimuli was observed in V1 (but not in V2 and V3) only for the unattended stimuli. Attention, on the other hand, was shown to correlate with an enhancement of the forward drive of information from V1 to V2 and V3, and was therefore proposed to reflect an increase in the postsynaptic gain of prediction error neurons (37). Expectation-attention interactions have also been demonstrated in several EEG studies focusing on ERP components occurring within 100-200ms after stimulus onset (38).

Critically, the methods used in the current study avoid some of the principal limitations of those studies:

1. The effects of expectation and attention were not confounded neither with each other nor by changes in low level properties of the sensory stimuli. This was accomplished by designing the paradigms of experiment 1 and Experiment 2 such that one cognitive domain was manipulated at a time by changing the behavioural task at hand.
2. The primary variable used in our analyses provides a direct neural measure of signal integration within the context of hierarchical processing. This was accomplished by using the Hierarchical Frequency Tagging method and by focusing analysis on IM components using the MSPCstim and MSPCres measures.

The above, with the combined results from multiple experiments, offer a unique advantage: the ability to obtain a direct and objective neurophysiological measure for the influence of expectation and attention on the integration of distinct streams of neural information in perception.

Instead of formulating the question at hand as whether expectation and attention increase or decrease neural activity, we place a spotlight on the role of signal integration in perception. In line with the predictive coding framework of perception, we view expectation and attention as distinct yet related mechanisms facilitating precision-weighted prediction error minimization, and serving the common goal of Bayes optimal perceptual inference. Our results highlight the role of feedback loops and integration of information across multiple hierarchical levels in the cortex, and relate expectation and attention to descending and ascending signals, influencing information integration at lower and higher cortical levels, respectively.

## Materials and Methods

### Hierarchical Frequency Tagging

The Hierarchical Frequency Tagging (HFT) method is based on the combination of two frequency tagging methods: a contrast modulation inducing Steady-State Visual Evoked Potential (SSVEP) tagging activity at lower visual areas, and an object-recognition modulation inducing Semantic Wavelet-Induced Frequency-Tagging (SWIFT) tagging activity at higher visual areas.

A detailed description of the method for creating SWIFT sequences can be found elsewhere (19). The crux of the SWIFT method is that it scrambles contours while conserving local low-level attributes such as luminance, contrast and spatial frequency. A SWIFT ‘sequence’ is a series of frames that, when presented sequentially, create a ‘movie’ that starts from a scrambled frame, progresses toward the middle (‘peak’) of the sequence where the original image can be briefly identified, after which the image becomes scrambled again. When such a SWIFT sequence is presented repetitively at a given frequency, the original images peaks at that frequency (i.e. once each SWIFT ‘cycle’) allowing the SWIFT tagging to be obtained (if the image is recognised).

To create a SWIFT sequence, we first scrambled the original image using the discrete Meyer wavelet and six decomposition levels. The local contour at each location and scale was then represented by a 3D vector. To create the sequence of frames, we then randomly selected two additional vectors of identical length and origin and defined the circular path that connected the three vectors (maintaining vector length along the path). We then performed additional cyclic wavelet-scrambling by rotating each original vector along the circular path and applying the inverse wavelet transform to obtain the image in the pixel domain. This way, we were able to smoothly scramble the original image by destroying contour information while conserving the local low-level attributes. The Matlab script for creating SWIFT sequences can be found at Koenig-Robert *et al*. (19).

In all experiments described here, we constructed each trial using SWIFT sequences created from one face and one house image, randomly selected for each trial from a pool of images. SWIFT sequences were presented consecutively, resulting in a ‘movie’ in which the original images (either the house or the face) were identifiable briefly around the peak of each such cycle (Video 1).

The SWIFT method preserves the low-level local visual properties across all frames within each sequence (cycle). These properties could differ significantly between a pair of SWIFT sequences used within a single trial (one for a face and the other for a house). Therefore, to preserve the low-level local visual properties across the whole *trial*, it was essential to alpha-blend (with equal weights) frames from both the house and the face sequences. As described below, a single SWIFT frequency was used in experiment 1 (Video 3), with either the face or the house image appearing in each cycle. To allow only one image to be recognised in each cycle, we created additional ‘noise’ sequences and alpha-blended them with the frames of the ‘image’ sequences presented in that cycle. This was done in the following way: First, we selected one of the scrambled frames from each of the original SWIFT sequences (the frame most distant from the original image presented at the peak of the cycle). Then, we created noise sequences by applying the SWIFT method on each of the selected scrambled frames. In this way, each original ‘image’ sequence had a corresponding ‘noise’ sequence that matched the low-level properties of the image sequence. Finally, during the actual presentation of the stimuli, we alpha-blended (with equal weights) the frames of the ‘image’ sequence that appeared in that cycle with the ‘noise’ frames of the other image category (see Figure 2A at Gordon *et al.* (16)). In Experiment 2, 15-30% of ‘image’ cycles were replaced with ‘noise’ cycles to make the counting task attentionally demanding (Video 4; see further details below). This way, the overall low level visual attributes were constant across all frames within the trial, regardless of the identifiable image in each cycle.

To minimise the possibility of confounding our results with the tagging of low-level visual features (specifically, by the cyclic repetition of the scrambled frames presented in between peaks) we created three sets of ‘image’ and ‘noise’ SWIFT sequence variants for each of the original images and continuously alternated between the sets during the trial. The timing of the transition between SWIFT variants (of a single original image) was designed to allow smooth transitions. This was done as follows: 1) If the image to be presented in the coming cycle is the same as the one presented in the current cycle (e.g. face-face), then the SWIFT variant of the ‘image’ sequence would be swapped at the peak of the current cycle (i.e. at the frame containing the original image). 2) If the image to be presented in the coming cycle is *not* the same as the one presented in the current cycle (e.g. face-house in experiment 1 or face-noise in Experiment 2), the transition between the image and its matching noise sequences would occur at the frame from which the ‘noise’ sequence was created. This way, we could better control for potential tagging of low-level visual features while ensuring smooth transitions between cycles.

For SSVEP, a global sinusoidal contrast modulation was applied on the whole movie. To avoid both excessively strong SSVEP signals and total blanks of the SWIFT sequences, the contrast modulation was limited to 70% of the full contrast range (from 30% to 100% of the original image contrast) (Video 2).

### Participants and experimental procedure

All participants were university students aged between 18 and 34 years. A total of n=15 out of 24 participants were included in the analysis of experiment 1, and n=11 out of 16 in experiment 2. Exclusion criteria were based on EEG quality and on task performance (see below.) In all experiments, participants were comfortably seated in a dimly lit room 55-60 cm in front of the monitor (LCD, 120Hz refresh rate). Stimuli were presented at the center of the screen over a grey background and participants were asked to keep their fixation at the center of the display. Participants were asked to minimise blinking or moving during each trial, but were encouraged to do so if needed in the breaks between trials. Trials always began by the participant pressing the spacebar.

Importantly, both new experiments performed in this study were designed such that the high-level factor was manipulated without modifying the stimuli used across experimental conditions. In other words, the predictability of the stimuli in the experiment 1 and the relevance of the stimuli in experiment 2 were manipulated through task instructions, without any confounds introduced by changes in low-level visual features.

### Behavioral tasks

#### Experiment 1-Expectation

Images appeared at 1.2Hz, alternating between a face and a house image in specific orders. In each trial, participants were required to perform one of two tasks such that each task was performed in half the trials.

In the image-repetition (IR) task, participants were asked to press the spacebar when they identified any image repeating itself either 3 or 4 times in a row. The specific number (3 or 4) was displayed on the screen prior to each trial. For example, for a trial in which the participant was instructed to look for an image repeating itself 3 times in a row, the face (F) and house (H) images may have been presented in the following order:

FHHFFHFHHFFHFHHFF**F**…

The ‘target’ image would then be the 18^th^ image (F shown in bold). Throughout the paper, these trials are referred to as the **image-repetition (IR)** trials.

In the pattern-violation (PV) task, participants were provided a pattern describing a series of 5-6 images *prior* to each trial, and were instructed to memorise the pattern well. Participants were told that the images presented in the trial would follow the given pattern, which will repeat itself over and over again. The task was then to press the spacebar as soon as they identified an image that violated the expected pattern.

For example, the following pattern may be verbally presented on the screen before a trial:

“Face, House, House, Face, Face, House”

After memorising the pattern, participants would begin the trial by pressing the spacebar. Images would then appear in the following order:

FHHFFH FHHFFH FHHFF**F**…

The ‘target’ image in this case would be the 18^th^ image shown in bold. the spaces between each pattern in the above example are provided here for illustration alone. In the actual trials, images appeared consecutively as per the 1.2 Hz SWIFT frequency. Throughout the paper, these trials are referred to as the **pattern-violation (PV)** trials.

As can be seen, the sequence of images and the target image are identical in both above examples, allowing differences in conscious perception, behavioural performance and evoked neural activity to be attributed strictly to the task-related requirements rather than the visual stimuli. We hoped that participants would not be aware of the underlying pattern when performing the IR task, rendering the series of images appear more random. As detailed in Results, this was indeed the case.

A global SSVEP contrast modulation was applied at 15Hz in all trials.

After 1-2 training trials, four 11-trial blocks were administered in the following order- PV block, IR Block, PV block, IR block. Every series that appeared in the PV block was repeated in the following IR block in a random order of trials, using a different set of face and house images. PV and IR trials were then analysed as paired.

Within 3 minutes from completing the experiment (i.e. after the end of the fourth block), participants were asked to compare the difficulty level between the PV and IR tasks, and to report whether they have noticed the underlying patterns in the IR tasks.

#### Experiment 2-Attention

HFT trials were constructed using house and face images. Contrary to experiment 1, here, the two image categories were presented at different SWIFT frequencies. In other words, the house and face images each cycled at its own frequency (0.8Hz and 1Hz, counterbalanced.) (Video 4). This allowed us to separately tag the EEG responses associated with the recognition of each image. Participants were instructed before each trial to count one of the image categories (e.g. ‘count houses’). Each image was therefore considered attended or unattended depending on what the participant was instructed to count during the trial. To ensure participants were actively paying attention to only one of the categories, rather than just following a certain ‘rhythm’, images were presented in only 70-85% of their respective cycles, and were substituted with their matching ‘noise’ sequence in the remaining cycles (Video 4). Trials were 31.5 seconds long, allowing the total amount of counted images to range from 17 to 26 images per trial. Participants were instructed not to expect trials with less than 10-15 images and were requested to be as accurate as possible when counting.

To further verify the dependence of the SWIFT response on attention, we added a third condition in which participants were required to perform a demanding central-attention task, leaving only minimal spatial attention elsewhere on the screen (39, 40). In this task, participants were instructed to attend to a central cross that varied in the height of the horizontal line (above or below the midline) and color (red, green or blue). The cross was updated at pseudo-random times, jittering over 850-1000ms to reduce frequency tagging by this stimulus (a constant time interval would yield an excessively strong tagging of the cross frequency). Participants were instructed to count the number of occurrences of two conjunction targets, defined as an upward and red cross or a downward and green cross. These central stimuli were also present in the count-face and count-house tasks, but they were irrelevant and ignored (Video 4).

Ten trials were administered in a random order for each of the three behavioural tasks (counting houses, faces or crosses), reaching 30 trials in total. The two images were presented at SWIFT frequencies of 0.8Hz and 1Hz, counterbalanced across trials. The global SSVEP contrast modulation was applied at 12Hz.

### Data acquisition and processing

Data were collected at two facilities, both using BrainProducts 64 scalp electrode EEG systems. Data for half of the participants from experiment 1 (n=12) and all participants from experiment 2 (n=16) were collected using an active-electrode actiCHamp system. Data for the other half of experiment 1 (n=12) were collected using a passive-electrode BrainAmp-MR system (*not* within an MR environment). Continuous EEG was sampled at 1,000 Hz for all participants.

Data processing was performed using the EEGLAB toolbox (41) in MATLAB. All data were resampled to 500 Hz. A high-pass filter was applied at 0.3 Hz and data was converted to average reference after replacing noisy electrodes. To define noisy electrodes, each sample point was regarded as being noisy if it was either greater than +80 μV (or lower than -80 μV), contained a sudden fluctuation greater than 30μV from the previous sample point, or if the signal was more than ±5 std from the mean of the trial data in each channel. Cycles in which over 2% of sample points were noisy were regarded as noisy cycles. Channels were replaced using spherical spline interpolation if they were considered noisy in over 10% of cycles. An additional CleanLine procedure was then applied to reduce AC power artifacts around 50Hz and 100Hz. The CleanLine plugin for EEGLAB (Mullen, 2012. Available online at: http://www.nitrc.org/projects/cleanline) reduces sinusoidal (line) noise while avoiding typical phase distortions that can be caused by notch filters.

### Exclusion criteria

Exclusion criteria were defined based on the quality of the EEG recording and the behavioral results. For the former, individual trials were marked for exclusion if over 10% of channels were considered noisy in that trial after pre-processing (as described above.) Participants with over 20% of bad trials were excluded from the analysis. A total of 5 participants from experiment 1 and 4 participants from experiment 2 were excluded based on this criteria for poor EEG recordings.

To ensure participants were sufficiently engaged with the tasks, we excluded participants whose responses were considered invalid in over 30% of trials. For experiment 1, responses were considered invalid if the spacebar was not pressed during the trial, or if it was pressed before the appearance of the target image (this criterion was applied on the PV trials). For Experiment 2, responses were considered invalid if they differed by more than +3 from the correct number (i.e. the actual number of times the attended image appeared in the trial). A total of 4 additional participants from experiment 1, and 1 additional participant from Experiment 2 were excluded based on this criteria for poor response accuracy.

### Spectral analysis

EEG amplitudes and phases were extracted at the tagging and intermodulation frequencies by applying the Fast Fourier Transform (FFT) over a predefined subset period in each trial. For experiment 2, the FFT was applied on the 25 second epoch ranging from 3-28 seconds from trial onset, yielding a half-bandwidth (i.e. frequency resolution) of 0.04Hz (=1/25s) (12,500 sample-points). For experiment 1 we applied the FFT on either 10 or 20 seconds of data, depending on the amount of data available in each trial (trial lengths in this experiment varied according to the location of the target image and the participant’s response). To reduce onset effects and the nosier signals often seen near trial onsets, we excluded the first SWIFT cycle of each trial from all analyses. Trials with more than 17 seconds of available data were zero-padded to 20 seconds and analysed as a 20-second trial, with a half-bandwidth of 0.05Hz (=1/20s). Trials with less than 17 seconds of data (but more than 10) were analysed using the first 10 seconds, with a half-bandwidth of 0.1Hz (=1/10s).

Signal-to-noise ratios (SNR) were computed by dividing the amplitude at any given frequency by the arithmetic mean amplitude across its neighbouring frequencies (42, 43). The specific number of neighbouring frequencies used for the SNR calculation depended on the length of data used in each analysis (as described above), ranging from 4 on each side (from f-0.4HZ to f+0.4HZ) for the 10-second epochs to 8 on each side (from f-0.32Hz to f+0.32Hz) for the 25-second epoch. Any neighbouring harmonic or IM frequency falling within that range was removed from the SNR calculation.

While a theoretically limitless number of IM components exist (all linear combinations of the fundamental input frequencies: n1f1 + n2f2, n = ±1,±2,±3…), we focused our primary analyses on the two lowest (2nd) order components (f1 ± f2) which tended to have the highest amplitude SNRs (as in Gordon *et al.*). Additional analyses were then performed on 3^rd^ and 4^th^ order IMs to broaden the scope of our investigation, as described in greater detail in Supplemental Information.

### Multi Spectral Phase Coherence

Distinct aspects of non-linear interactions may be revealed by examining both phase and amplitude information. Applying phase analyses rather than amplitude analyses alone may have several advantages. First, amplitude and phase information may indicate different aspects of neural processing, with the phase-coherence believed to reflect the relative timing of neural activity (44). Second, noise that is not associated with stimulus processing is by definition not time-locked to stimulus onset, and is therefore not expected to demonstrate any phase consistency across trials. Consequently, phase analyses may be more robust to noise, potentially allowing the detection of genuine response components even when the amplitude is low.

The phase-coupling measure we use in this study is the multi-spectral phase coherence (MSPC) introduced by Yang *et al.* (28). The MSPC is especially useful for the study of intermodulation components since rather than comparing a frequency phase to a time-locked event, it allows it to be compared against the phases of the fundamental frequencies. In that sense, instead of asking ‘how strongly is the IM phase driven by a given event in time’ (as with the classic phase-locking factor), the MSPC asks ‘how strongly is the IM phase determined by the phases of the input frequencies driving the IM response’. The MSPC can be calculated between input (I) frequencies f1,f2…fr and any given output (O) IM (or harmonic) frequency 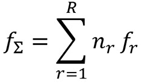 (where *f*Σ is a linear combination of the input frequencies with weights n1,n2…nr). Mathematically, this is given by the formula:

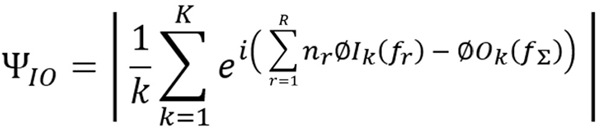

Where Ø*I_k_*(*f_r_*) is the phase of the input frequency f_r_ (in epoch k), and Ø*O_k_*(*f*_Σ_) is the phase of the output IM (in that same epoch).

There are various additional conceptual differences between the more commonly used phase-locking value (PLV) and the MSPC measure we apply here. First, for PLV, the phase of only one frequency of interest is extracted from each epoch to calculate PLV values. Here, for each MSPC calculation, we extract the phases of three frequencies from each epoch-two fundamental and one IM component phase. Second, PLV is often used to examine the coherence between different neural signals by testing the consistency of the difference (Δϕ) between phases of distant channels. Here, we test the consistency of the difference between the phases of the fundamental frequencies and the IM component within a channel.

The logic behind performing within-channel calculations is that the sources associated with the SWIFT, the SSVEP and the IM responses should, in principle, have an additive (and therefore separable) influence on the resulting EEG signal at each channel. Moreover, if we assume that for any given channel, the phase of the signal associated with the activity of a particular source depends on a characteristic time delay, then the dynamics of a single electrode should contain information about the dynamics of multiple electrodes (45). Therefore, we used within-channel data to examine the relationship between the IM and the SWIFT and SSVEP phases while avoiding the complexity involved in cross-channel computations.

A novel distinction we introduce here is between the calculation of the MSPC based on the stimulus and the response. We define the stimulus phase as the phase of the SSVEP contrast modulation and the phase of the SWIFT sequence relative to the original image. We define the response phase as the phases of the Fourier Transform of the EEG at the relevant frequencies (i.e., the SWIFT and SSVEP frequencies). MSPC values were calculated for each channel individually.

We reasoned that separately considering the stimulus phase and the response phase as the driving ‘inputs’ of the IM response may allow a distinction between interactions occurring at lower and higher levels of hierarchical processing, respectively. The logic behind this is as follows: While the stimulus phases should relate strongly to the phases of low-level retinal signals, the response phases reflect activity occurring further up the visual pathway. Importantly however, the primary sources of the SSVEP and SWIFT EEG signals are not the same but rather low and high visual levels, respectively (19). If the processing time-delays between retinal input and activity at SSVEP-generating regions and SWIFT-generating regions were different, yet *constant*, the MSPC measure should not differ when calculated based on the stimulus or based on the response phases of the SSVEP and SWIFT. This is because MSPC measures the degree of consistency between an IM frequency phase and the combined SSVEP and SWIFT phases across epochs, which should not depend on the addition of constants. However, the processing times leading to activity at SSVEP- and SWIFT-generating regions may not be constant across all stimuli. For example, the timing can vary depending on factors such as expectation, attention, image visibility, recognition, etc. Therefore, when examined across multiple epochs, the IM response phase may have a different relationship with the combination of the SSVEP and SWIFT *stimulus* phases than with the combination of the SSVEP and SWIFT *response* phases. This is the logic behind our assumption that these two measures, which we denote MSPCstim and MSPCres, may reveal different information about interactions occurring at lower and at higher levels of the visual hierarchy, respectively.

MSPC measures were calculated in a within-trial manner by dividing each trial into a series of 5-second (experiment 1) or 10-second (experiment 2) epochs, with a 1-second step from epoch to epoch (shorter epochs were used for experiment 1 since the average length of data available for each trial was shorter than those of experiment 2).

### Expectation-Attention interaction (Reanalysis of eLife)

In addition to the two experiments described above, new analyses were performed on the data published in Gordon *et al.* (16). A detailed description of the paradigm and analysis methods can be found in the original publication. In brief, house and face SWIFT cycles were presented in each trial in a pseudo-random order and participants were asked to count either the houses or the faces. Certainty (expectation) levels were categorised based on the proportion of house and face images appearing in each trial, and ranged from low certainty (faces and houses appeared at nearly 50% of cycles each) to high certainty (one of the two images appeared in nearly 100% of cycles).

Importantly, while the behavioural task allowed us to verify participants were engaged with the task, it introduced a within-trial difference between the attended (counted) and unattended images, which was not analysed in that study. Given that one underlying SWIFT frequency was used (each cycle peaking at either the face or the house image) and the spectral analysis was performed on full trials, face- and house-driven EEG responses could not be distinguished from each other in the frequency domain. The effect of expectation on amplitude SNRs was examined in the original study using the linear mixed-effects (LME) analyses with a model that included certainty as the fixed effect and channels nested within participants as random effects.

Here, we first calculated new MSPCstim and MSPCres measures for the 2^nd^ order IM components. Then, to allow the additional examination of potential interactions between expectation and attention (counted vs. uncounted images), an additional attention-dependent variable was added- the attentional category of the more frequent (higher-certainty) image. In other words, this variable indicated whether the image presented in most cycles (above half) in any given trial was the counted or the not-counted image. This way, we could now distinguish between the effects of high expectation for attended vs for unattended images. In the new LME model, expectation, attention, and an expectation-attention interaction were included as the fixed effects. Random effects included a random intercept for frequency nested within channels nested within participants, and random expectation, attention and interaction slopes for each participant.

To examine the consistency of the full interaction LME model with the results from experiment 1 and Experiment 2, we performed additional post-hoc tests to individually examine expectation and attention. For conditions similar to those of experiment 1, we first tested the effect of expectation within the attended condition. Then, for conditions similar to those of Experiment 2, we used a median split to reduce expectation to two bins (expected and unexpected) and we tested the effect of attention within the expected condition.

As in the original study, we tested for the significance of a given factor or interaction by performing a likelihood ratio test between the full model, as described above, and the reduced model which did not include the factor in question (46).

## Acknowledgments

We would like to thank Ms. Kavya Raj for her technical assistance at the early stages of this study.

This study was supported by funding from Australian Research Council FT120100619, DP180100396, DP180104128 Naotsugu Tsuchiya, DP160102770 Jakob Hohwy, Templeton World Charity Foundation, Inc (TWCF0199) Naotsugu Tsuchiya.

## Supplemental Information

### Spectra of experiment 1 and experiment 2

**Figure S1.**
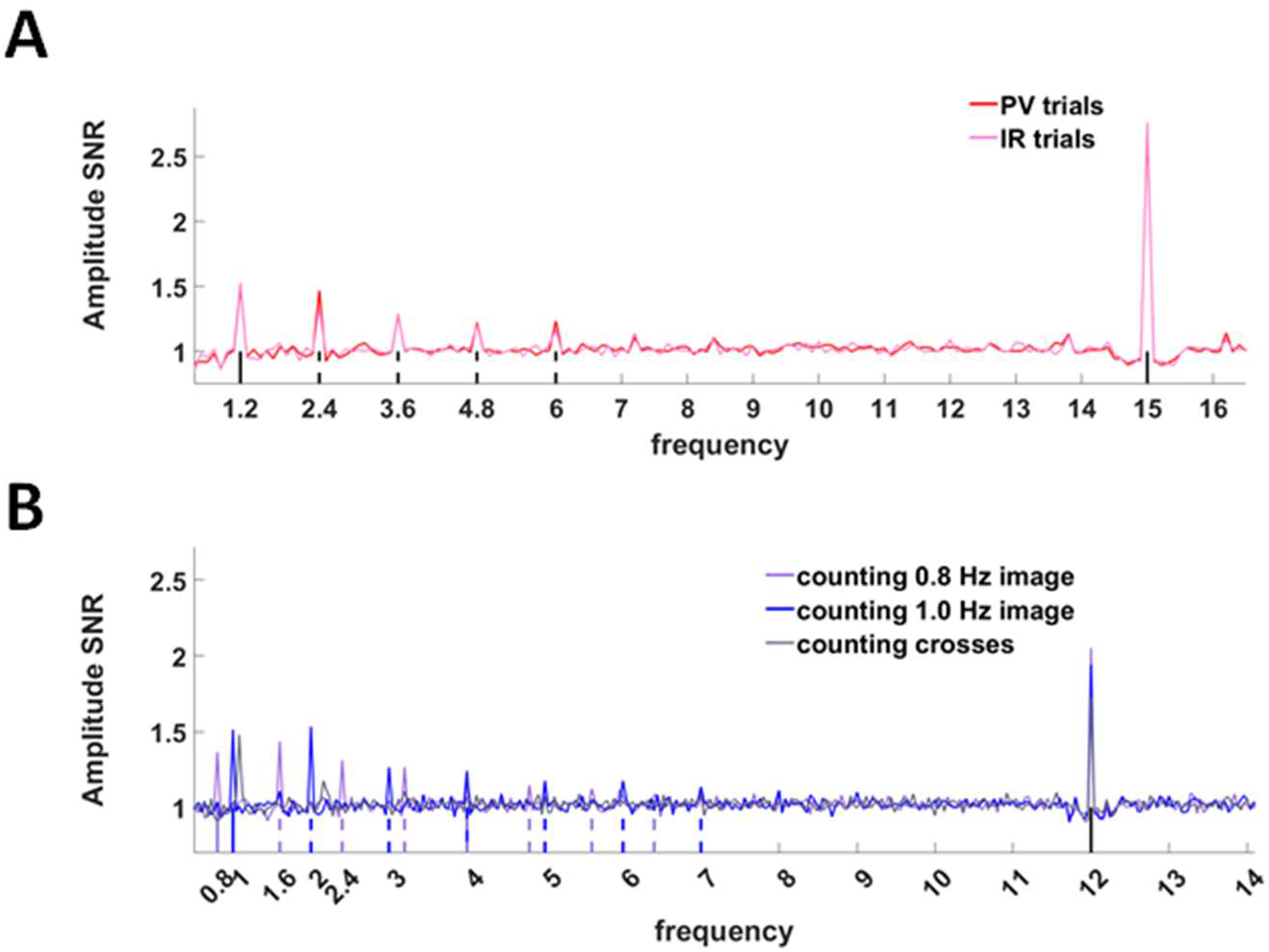
Related to Figure 3 and Figure 4 – Amplitude SNRs demonstrate successful frequency tagging in Experiments 1 and 2. Results of the fast Fourier transform (FFT) averaged across all electrodes, trials and participants. Amplitude SNR peaks can be seen at the tagging frequencies (solid lines) and their harmonics (dashed lines) in both experiment 1 (A) (SWIFT: 1.2 Hz and SSVEP: 15 Hz, N=15) and Experiment 2 (B) (two SWIFT: 0.8Hz and 1Hz, and SSVEP: 12 Hz, N=11). Note that in Experiment 2 no SWIFT tagging was obtained when counting crosses (the peak at ~1.08 Hz matches the average amount of time between cross presentations which was 925ms).

### MSPC and Amplitude measures

We performed a series of analyses to examine the potential confound between the MSPC and amplitude measures. Specifically, given the significantly reduced SWIFT amplitude tagging for unattended images (Figure S1), we wished to examine whether the reduction of MSPCres in the unattended images was a consequence of a potential decrease in the reliability of phase measures when amplitudes are low. For this, we performed additional analyses to evaluate the relation between MSPCres and the IM amplitude, and between MSPCres and the SWIFT amplitude (note that while different SWIFT frequencies were used in Experiment 2 for the attended and unattended images, only a single SSVEP frequency was applied, hence the SSVEP cannot be differentiated for the attended and unattended images in that experiment).

We performed all correlation analyses using the Spearman correlation within each participant. This way, we could avoid unwanted effects that can result from differences in the EEG topographies across participants or from the potentially different distributions of the MSPCres and amplitude measures. To do so, we first computed the MSPCres and the amplitude SNR for the second order IMs (f1+-f2) and the SWIFT frequency across trials for a given EEG channel for each participant. Then, for each participant we calculated the Spearman correlation between the MSPCres and the amplitude measures across all electrodes.

To examine the relationship between the MSPCres and the amplitude SNR of the *IMs* we performed two tests: 1) a correlation analyses between MSPCres and the amplitude SNR of the 2^nd^ order IMs, and 2) a LME model (like the one used in Figure 5) with amplitude SNR as the predicted variable instead of the MSPCres. The results of the within-participant correlations analyses demonstrated low correlations between the MSPCres and the IM amplitude SNRs (average R^2^=0.06, min R^2^=0.00 and max R^2^=0.22 across all participants). The results of the LME indicated that while MSPCres decreased with expectation in the unattended condition, the amplitude SNR in fact trended upward.

We also computed correlation between the MSPCres and the amplitude SNR of the *SWIFT* frequency. Also here, the within-participant correlations demonstrated low correlations (average R^2^=0.04, min R^2^=0.00 and max R^2^=0.19 across all participants).

Taken together, these results indicated that the amplitude of the IM and SWIFT frequencies alone cannot account well for the behaviour of the MSPCres.

### Interaction of attention and expectation-Related to Figure 5

After differentiating expectation and attention in Experiments 1 and 2, and establishing that both expectation and attention are associated with enhanced IMs, we returned to our previously published data (16) to examine the interaction between these factors, and to evaluate the consistency of the results from that study with those of Experiments 1 and 2 presented here.

In Gordon *et al.*, we analyzed frequency-domain amplitudes for SSVEP, SWIFT and IM, as a function of the number of counted stimuli, which confounded the effects of expectation and attention. Here, we separated the main effects of expectation and attention from their interaction within that single paradigm, using the MSPC-based measures as above. To do so, an additional ‘attention’ variable was added to the model, allowing us to examine the interaction between expectation and attention. The new ‘attention’ variable indicated, for each trial, whether it was the counted (attended) or the uncounted image that appeared in most cycles. Then, a likelihood ratio test between was performed between the full LME interaction model (including expectation, attention and the expectation-attention interaction as the fixed effects) and the reduced model (including expectation and attention as fixed effects, without the interaction term) (see Methods).

The interaction between expectation and attention was not significant for MSPCstim (χ2 = 3.47, p>0.05) but it was indeed highly significant for MSPCres (χ2 = 19.56, p<0.001). In fact, the slope of MSPCres against expectation was negative for unattended images (χ2 = 5.05, p<0.05) (Figure 5). These results are interpreted further in Discussion.

To examine the consistency of the results from this data with those of experiment 1 and 2, we performed individual post-hoc analyses for expectation and attention. For conditions more similar to those of experiment 1, we tested the effect of expectation within the attended condition. For conditions more similar to those of Experiment 2, we used a median split to reduce expectation into two bins (expected and unexpected) and we then tested the effect of attention within the high expectation bin. Consistent with the results from Experiments 1, the effect of expectation within the attended condition was significant for MSPCstim (χ2 = 7.35, p<0.01) but not for MSPCres (χ2 = 1.86, p>0.05), and the effect of attention within the expected condition was more notable for MSPCres (χ2 = 12.11, p<0.001) than for MSPCstim (χ2 = 4.53, p=0.033). The effect of attention on MSPCstim did not survive the false discovery rate (FDR) correction for multiple comparisons (p= 0.066 after FDR adjustment).

### Higher order IMs

After establishing the modulatory influence of expectation and attention on the 2^nd^ order IMs (f2±f1), we performed additional analyses on the 3rd (f2±2f1, 2f2±1f1) and 4th (2f2±2f1) order components. High-order IMs can arise from a sequence of lower-order computations. Here, we examined MSPCres at the higher-order IMs to compare between two plausible computational sequences.

Before doing so, we first examined whether the effects of the behavioural manipulations were evident in the 3^rd^ and 4^th^ order IMs. For experiment 1 (expectation modulation), only the 4th order IMs demonstrated similar modulatory influence of expectation, with MSPCstim of those IMs being higher in the PV (expected) compared to the IR (unexpected) trials (χ2= 8.04, p<0.01). This effect was nevertheless not as significant as the effect observed for the 2nd order components (χ2= 22.9, p<0.001). For experiment 2 (attention modulation), both the 3rd and the 4th order IMs demonstrated a similar modulatory influence of attention, with MSPCres being significantly higher for counted (attended) compared to non-counted (unattended) images (χ2 > 20 and p≪0.001 for both comparisons).

The 4th order IMs can be described as the harmonic of the 2^nd^ order IM (i.e. F2,F1 → (f2+f1) → 2(f2+f1)), or, alternatively, as the 2^nd^ order IM between the harmonics of the fundamental frequencies (i.e. F2,F1 → 2f2, 2f1 → 2f2+2f1). To distinguish between these two possibilities, we ran additional MSPCres analyses in which we defined either the IM components f1+f2 and f1-f2, or the harmonics 2f2 and 2f1 as the driving input frequencies of the 4^th^ order 2f2±2f1 IMs. In other words, we examined whether the 4th order IMs reflects ‘early-interaction’, driven by the 2^nd^ order IMs (as would be the case if an initial interaction between the input signals is followed by another non-linear process) or ‘late-interaction’, driven by the harmonics of the fundamental frequencies (as would be the case if the input signals are processed individually and then interact).

For statistical analysis, we defined an LME model in which attention, the MSPC computation method (i.e. bases on 2f1 and 2f2 or based on f2-f1 and f2+f1) and the interaction between the two were included as the fixed effects. Random effects included a random intercept for frequency nested within channels nested within participants, and random attention, MSPC computation method and interaction slopes for each participant.

The MSPCres of the 4th order IMs was significantly higher when calculated based on the harmonics of the fundamental frequencies (2f1 and 2f2) than when calculated based on their 2^nd^ order IMs (f2+f1 and f2-f1) (χ2= 27.2, p<0.001) (Figure S2). These results therefore favour the ‘late-interaction’ option in which the input signals are processed individually before interacting with each other. Indeed, the interaction between attention and the MSPC computation method was significant (χ2= 8.4, p<0.01), indicating that attention had a significantly greater influence on the degree to which the 4th order IMs were driven by the 2f1 and 2f2 harmonics than by the f1+f2 IMs. These results provide additional support linking the MSPCres measure and the attentional modulation to interactions occurring at later stages than where initial stimulus processing and interactions occurs.

**Figure S2.**
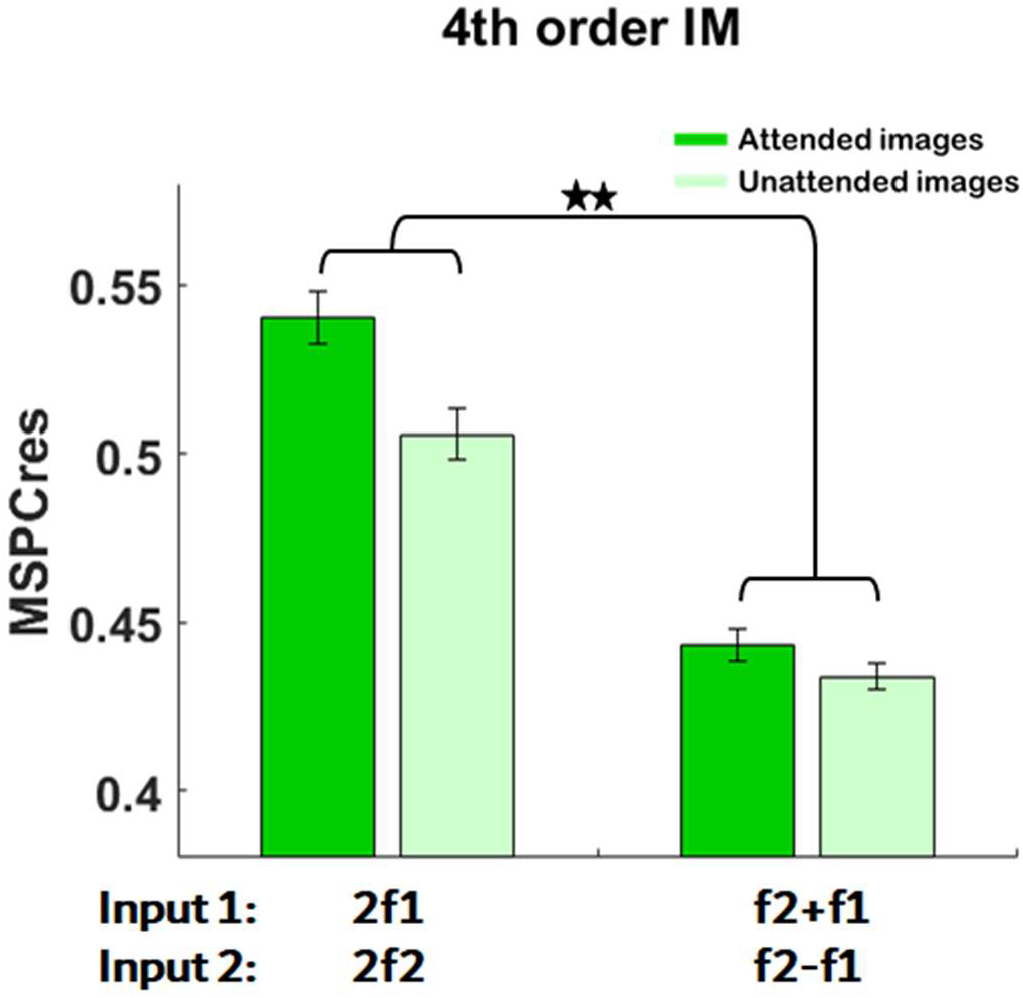
Related to Figure 4 – The 4^th^ order IMs reflect the late interaction (i.e., the second order IM between the second harmonics) rather than the early interaction (i.e., the second harmonics of the second order IM) Additional analyses of the 4^th^ order IMs were performed to compare between two potential two-stage 2^nd^ order sequences-non-linear processing of each of the input signal followed by an interaction between the two (i.e. F2,F1 → 2f2, 2f1 → 2f2+2f1, leŌ bars), or an interaction between the input signals followed by an additional non-linear process (i.e. F2,F1 → (f2+f1) → 2(f2+f1), right bars). Higher MSPCres values were obtained, for both the attended and the unattended images, when defining the 2^nd^ harmonics of the SWIFT and SSVEP response frequencies as the driving input signals. This indicates that the 4^th^ order IMs are driven more by these 2^nd^ harmonics than by their 2^nd^ order IMs. Furthermore, attention had a significantly greater influence on the degree to which the 4th order IMs were driven by the 2f1 and 2f2 harmonics than by the f1+f2 IMs. These results are consistent with the notion that the attention modulation influences processes occurring at later stages than where initial input processing and interactions occur.

## References

1. Doya K. Bayesian brain: probabilistic approaches to neural coding. Cambridge, Mass.: MIT Press; 2007. 326 p. p.

2. Friston K. A theory of cortical responses. Philosophical transactions of the Royal Society of London Series B, Biological sciences. 2005;360(1456):815–36.

3. Feldman H, Friston KJ. Attention, Uncertainty, and Free-Energy. Frontiers in Human Neuroscience. 2010;4:215.

4. Kersten D, Mamassian P, Yuille A. Object perception as Bayesian inference. Annual review of psychology. 2004;55:271–304.

5. Lee TS, Mumford D. Hierarchical Bayesian inference in the visual cortex. J Opt Soc Am A Opt Image Sci Vis. 2003;20(7):1434–48.

6. Schroger E, Marzecova A, SanMiguel I. Attention and prediction in human audition: a lesson from cognitive psychophysiology. Eur J Neurosci. 2015;41(5):641–64.

7. Summerfield C, de Lange FP. Expectation in perceptual decision making: neural and computational mechanisms. Nat Rev Neurosci. 2014;15(11):745–56.

8. Summerfield C, Egner T. Expectation (and attention) in visual cognition. Trends Cogn Sci. 2009;13(9):403–9.

9. Todorovic A, Schoffelen JM, van Ede F, Maris E, de Lange FP. Temporal expectation and attention jointly modulate auditory oscillatory activity in the beta band. PLoS One. 2015;10(3):e0120288.

10. Robinson JE, Breakspear M, Young AW, Johnston PJ. Dose-dependent modulation of the visually evoked N1/N170 by perceptual surprise: a clear demonstration of prediction-error signalling. Eur J Neurosci. 2018.

11. Rao RPN, Ballard DH. Predictive coding in the visual cortex: a functional interpretation of some extra-classical receptive-field effects. Nature neuroscience. 1999;2(1):79–87.

12. Kok P, Rahnev D, Jehee JF, Lau HC, de Lange FP. Attention reverses the effect of prediction in silencing sensory signals. Cereb Cortex. 2012;22(9):2197–206.

13. Garrido MI, Rowe EG, Halasz V, Mattingley JB. Bayesian Mapping Reveals That Attention Boosts Neural Responses to Predicted and Unpredicted Stimuli. Cereb Cortex. 2018;28(5):1771–82.

14. Auksztulewicz R, Friston K. Repetition suppression and its contextual determinants in predictive coding. Cortex. 2016;80:125–40.

15. Friston K, Kiebel S. Predictive coding under the free-energy principle. Philosophical transactions of the Royal Society of London Series B, Biological sciences. 2009;364(1521):1211–21.

16. Gordon N, Koenig-Robert R, Tsuchiya N, van Boxtel JJ, Hohwy J. Neural markers of predictive coding under perceptual uncertainty revealed with Hierarchical Frequency Tagging. Elife. 2017;6.

17. Norcia AM, Appelbaum LG, Ales JM, Cottereau BR, Rossion B. The steady-state visual evoked potential in vision research: A review. J Vis. 2015;15(6):4.

18. Vialatte FB, Maurice M, Dauwels J, Cichocki A. Steady-state visually evoked potentials: focus on essential paradigms and future perspectives. Prog Neurobiol. 2010;90(4):418–38.

19. Koenig-Robert R, VanRullen R. SWIFT: a novel method to track the neural correlates of recognition. NeuroImage. 2013;81:273–82.

20. Di Russo F, Pitzalis S, Stella A, Spinelli D, Hillyard SA. Identification of the cortical sources of the steady-state visual evoked potential: A vep-fMRI co-registration study. Journal of Vision. 2006;6(6):1095.

21. Koenig-Robert R, VanRullen R, Tsuchiya N. Semantic Wavelet-Induced Frequency-Tagging (SWIFT) Periodically Activates Category Selective Areas While Steadily Activating Early Visual Areas. PLoS ONE. 2015;10(12):e0144858.

22. Connor CE, Brincat SL, Pasupathy A. Transformation of shape information in the ventral pathway. Curr Opin Neurobiol. 2007;17(2):140–7.

23. DiCarlo JJ, Zoccolan D, Rust NC. How does the brain solve visual object recognition? Neuron. 2012;73(3):415–34.

24. Regan D. Evoked potential studies of visual perception. Can J Psychol. 1981;35(2):77–112.

25. Boremanse A, Norcia AM, Rossion B. An objective signature for visual binding of face parts in the human brain. Journal of Vision. 2013;13(11):6-.

26. Gundlach C, Muller MM. Perception of illusory contours forms intermodulation responses of steady state visual evoked potentials as a neural signature of spatial integration. Biological psychology. 2013;94(1):55–60.

27. Zhang P, Jamison K, Engel S, He B, He S. Binocular rivalry requires visual attention. Neuron. 2011;71(2):362–9.

28. Yang Y, Solis-Escalante T, Yao J, Daffertshofer A, Schouten AC, van der Helm FCT. A General Approach for Quantifying Nonlinear Connectivity in the Nervous System Based on Phase Coupling. International Journal of Neural Systems. 2016;26(01):1550031.

29. McMains SA, Fehd HM, Emmanouil TA, Kastner S. Mechanisms of feature- and space-based attention: response modulation and baseline increases. J Neurophysiol. 2007;98(4):2110–21.

30. Hohwy J. Attention and Conscious Perception in the Hypothesis Testing Brain. Frontiers in psychology. 2012;3:96.

31. Kanai R, Komura Y, Shipp S, Friston K. Cerebral hierarchies: predictive processing, precision and the pulvinar. Philosophical transactions of the Royal Society of London Series B, Biological sciences. 2015;370(1668).

33. den Ouden HE, Friston KJ, Daw ND, McIntosh AR, Stephan KE. A dual role for prediction error in associative learning. Cereb Cortex. 2009;19(5):1175–85.

32. Kirschstein T, Kohling R. What is the source of the EEG? Clin EEG Neurosci. 2009;40(3):146–9.

34. Kok P, Jehee Janneke FM, de Lange Floris P. Less Is More: Expectation Sharpens Representations in the Primary Visual Cortex. Neuron. 2012;75(2):265–70.

35. Murray MM, Wylie GR, Higgins BA, Javitt DC, Schroeder CE, Foxe JJ. The spatiotemporal dynamics of illusory contour processing: combined high-density electrical mapping, source analysis, and functional magnetic resonance imaging. The Journal of neuroscience: the official journal of the Society for Neuroscience. 2002;22(12):5055–73.

36. Jiang J, Summerfield C, Egner T. Attention sharpens the distinction between expected and unexpected percepts in the visual brain. The Journal of neuroscience: the official journal of the Society for Neuroscience. 2013;33(47):18438–47.

37. Kok P, Rahnev D, Jehee JFM, Lau HC, de Lange FP. Attention Reverses the Effect of Prediction in Silencing Sensory Signals. Cerebral Cortex. 2012;22(9):2197–206.

38. Hsu YF, Hamalainen JA, Waszak F. Both attention and prediction are necessary for adaptive neuronal tuning in sensory processing. Front Hum Neurosci. 2014;8:152.

39. Bahrami B, Lavie N, Rees G. Attentional load modulates responses of human primary visual cortex to invisible stimuli. Curr Biol. 2007;17(6):509–13.

40. van Boxtel JJ, Tsuchiya N, Koch C. Opposing effects of attention and consciousness on afterimages. Proceedings of the National Academy of Sciences of the United States of America. 2010;107(19):8883–8.

41. Delorme A, Makeig S. EEGLAB: an open source toolbox for analysis of single-trial EEG dynamics including independent component analysis. Journal of Neuroscience Methods. 2004;134(1):9–21.

42. Srinivasan R, Russell DP, Edelman GM, Tononi G. Increased Synchronization of Neuromagnetic Responses during Conscious Perception. The Journal of Neuroscience. 1999;19(13):5435–48.

43. Tononi G, Edelman GM. Consciousness and Complexity. Science. 1998;282(5395):1846–51.

44. Sauseng P, Klimesch W. What does phase information of oscillatory brain activity tell us about cognitive processes? Neurosci Biobehav Rev. 2008;32(5):1001–13.

45. Tajima S, Yanagawa T, Fujii N, Toyoizumi T. Untangling Brain-Wide Dynamics in Consciousness by Cross-Embedding. PLoS Comput Biol. 2015;11(11):e1004537.

46. Bates D, Mächler M, Bolker B, Walker S. Fitting Linear Mixed-Effects Models Using lme4. 2015. 2015;67(1):48.

